# Pollen morphology, deep learning, phylogenetics, and the evolution of environmental adaptations in *Podocarpus*

**DOI:** 10.1101/2025.01.07.631571

**Authors:** Marc-Élie Adaimé, Michael A. Urban, Shu Kong, Carlos Jaramillo, Surangi W. Punyasena

## Abstract

- *Podocarpus* pollen morphology is shaped by both phylogenetic history and the environment. We analyzed the relationship between pollen traits quantified using deep learning and environmental factors within a comparative phylogenetic framework.
- We investigated the influence of mean annual temperature, annual precipitation, altitude, and solar radiation in driving morphological change. We used trait-environment regression models to infer the temperature tolerances of 31 Neotropical *Podocarpidites* fossils. Ancestral state reconstructions were applied to the *Podocarpus* phylogeny with and without the inclusion of fossils.
- Our results show that temperature and solar radiation influence pollen morphology, with thermal stress driving an increase in pollen size and higher UV-B radiation selecting for thicker corpus walls. Fossil temperature tolerances inferred from trait-environment models aligned with paleotemperature estimates from global paleoclimate models. Incorporating fossils into ancestral state reconstructions revealed that early ancestral *Podocarpus* lineages were likely adapted to warm climates, with cool-temperature tolerance evolving independently in high-latitude and high-altitude species.
- Our results highlight the importance of deep learning-derived features in advancing our understanding of plant environmental adaptations over evolutionary timescales. Deep learning allows us to quantify subtle interspecific differences in pollen morphology and link these traits to environmental preferences through statistical and phylogenetic analyses.

## Introduction

Plant phenotypic traits are shaped by the environment over evolutionary timescales. Morphology can therefore provide insight into how plant species adapted to long-term climatic and environmental changes. Among these phenotypic traits, pollen morphology holds particular importance due to its role in reproduction and the strong selective pressures associated with pollination biology (Fatmi *et al*., 2020). The production and morphology of pollen grains are critical to reproductive success in conifers (Owens *et al*., 1998; Leslie, 2010; Leslie *et al*., 2015). In angiosperms, studies of Myrtales and Asterlales have shown that pollen morphology reflects both phylogenetic history and environmental pressures (Kriebel *et al*., 2017; Jardine *et al*., 2022), making it a useful metric for understanding how plants have adapted to diverse habitats over time.

Characterizing pollen morphology and quantifying its variability, however, presents a significant challenge. Differences in pollen shape and texture are often described qualitatively, using the rich descriptive terminology of palynology (Punt *et al*., 2007). In saccate pollen grains, quantitative features include grain size, saccus height, width, and shape, angle of saccus attachment, endoreticulate patterning of the sacci, the relative size of sacci to corpus, the width of the saccus at the point of attachment, and the thickness of the corpus wall (Punyasena *et al*., 2012; Masetto & Lorscheitter, 2016; Zavattieri *et al*., 2018). In many clades, including *Podocarpus*, pollen morphology tends to be highly conserved (Pocknall, 1981; Ledru *et al*., 2007; Adaimé *et al*., 2024a), suggesting that much of the morphological variation within pollen is constrained by phylogenetic history. However, subtle morphological differences within these constraints potentially reflect adaptive responses to environmental stressors. Traditional morphometric methods often fail to capture these fine-scale morphological differences, leaving important gaps in our understanding of how environmental factors shape pollen evolution across lineages.

Advances in deep learning, however, provide powerful tools for representing morphology. Convolutional neural networks (CNNs) can quantify high-level, abstract morphological features that are difficult to discern using conventional approaches (Adaimé *et al*., 2024a,b), offering insights for phylogenetic applications (Adaimé *et al*., 2024a,b). These numeric characterizations allow us to statistically isolate the features of pollen morphology that are directly shaped by environmental pressure and are less constrained by phylogenetic history. This enables us to model the relationship between environmental factors and complex morphological characters in pollen grains. Using phylogenetic comparative methods, we can now extract and interpret these features in an evolutionary context, identifying key adaptations and revealing how environmental pressures have driven morphological diversification in pollen over time.

The abundance of pollen in the fossil record allows high-resolution tracking of historical environmental influences on plant evolution, capturing phenotypic shifts and adaptations across different lineages. But despite its ubiquity, few studies have explored how pollen traits have evolved in response to environmental pressures. Previous research has linked extant pollen morphology to environmental factors (e.g, UV-A and UV-B exposure; Yeloff *et al*., 2008), but these studies often lack an evolutionary framework, leaving gaps in our understanding of how traits like exine thickness or pollen grain structure have historically evolved along a phylogeny. Studies using fossil pollen have examined traits such as pollen size in relation to past temperature or moisture conditions (e.g., Ejsmond *et al*., 2011, 2015; Griener & Warny, 2015; McCulloch *et al*., 2022), but have mainly focused on paleoecological applications. Traits are used as proxies to reconstruct past climate change without explicitly mapping their evolution over time. Previous studies have also predominantly focused on more easily quantifiable traits such as grain size or exine thickness, leaving finer details like textural patterning or subtle differences in the grain’s internal structure largely unexamined.

*Podocarpus*, the largest and most widespread genus in the Podocarpaceae, spans a broad range of climatic zones (Figs. 1 and 2), from humid tropics to cold, high-altitude regions (Punyasena *et al*., 2011; Quiroga *et al*., 2016). This environmental diversity makes *Podocarpus* an ideal clade for investigating how species adapt to varying climatic conditions through the study of pollen morphology. Although highly conserved (Pocknall, 1981; Ledru *et al*., 2007), *Podocarpus* pollen morphology encodes much more evolutionary information than had been previously recognized (Adaimé *et al*., 2024a), providing an opportunity to examine species-specific trait-environment relationships. The fossil pollen morphotype *Podocarpidites* is the palynological form genus for *Podocarpus* and has a global distribution from the Late Cretaceous through the Cenozoic.

**Figure 1.**
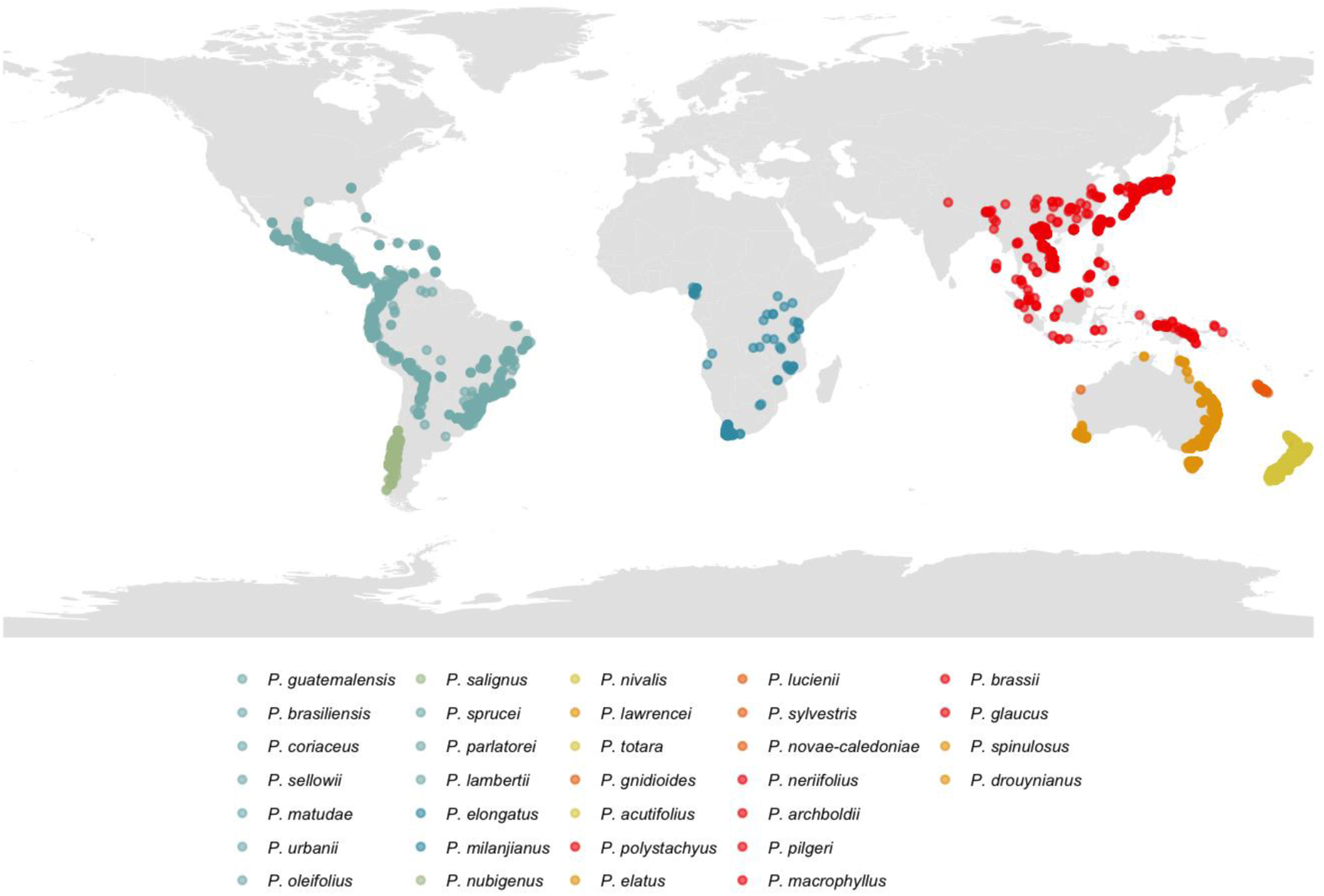
Global distribution of the 32 extant *Podocarpus* species including in our phylogenetic analyses. Species occurrences are based on data obtained from the Global Biodiversity Information Facility (GBIF). Occurrences were filtered by plotting the centroids of each species’ geographic distribution and removing records that were located more than 7,000 km from their respective centroids.

**Figure 2.**
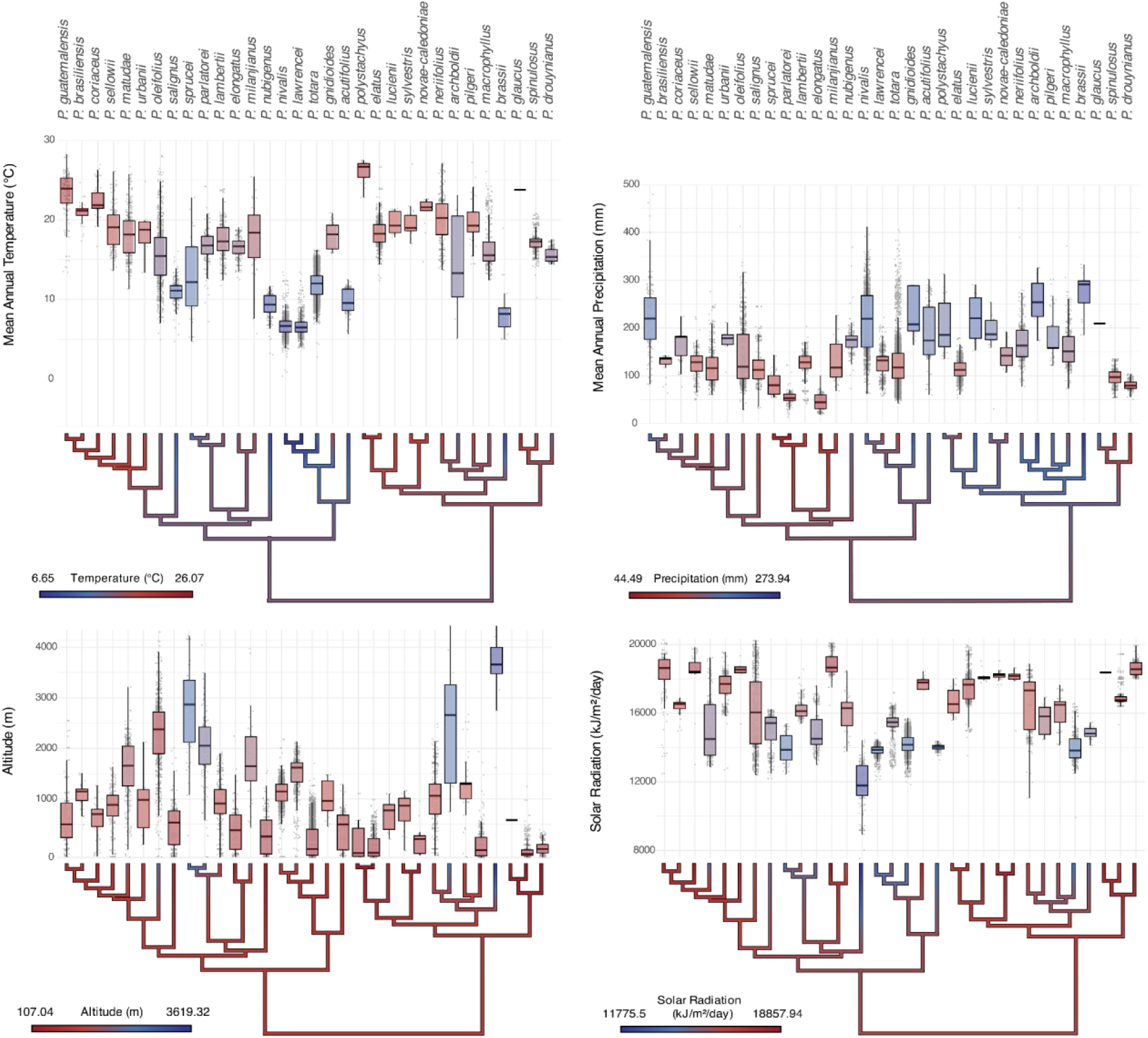
Environmental tolerances of the 32 extant *Podocarpus* species and their most likely ancestral states along the phylogeny. The box plots display the species’ environmental tolerances (mean annual temperature, mean annual precipitation, altitude, and solar radiation) extracted from WordClim 2 (Fick & Hijmans, 2017). The phylogenetic trees beneath each box plot show the reconstructed ancestral states for these variables using maximum likelihood inference under an Ornstein-Uhlenbeck (OU) model (α = 0.1). The gradient color bars along the trees represent the predicted values of environmental tolerances for both extant and ancestral nodes, with warmer and cooler colors indicating higher and lower values, respectively.

In this study, we used abstract morphological features derived from CNNs and phylogenetic comparative methods to test hypotheses of pollen evolution in response to environmental factors. Our results indicate that temperature and solar radiation most strongly influence *Podocarpus* pollen morphology, independent of shared evolutionary history. We used phylogenetically constrained trait-environment regression models to infer the environmental tolerances of 31 fossil *Podocarpidites* specimens from South America, spanning the Late Cretaceous to the Pliocene, and compared these to their predicted paleotemperature preferences, based on the specimens’ ages and paleogeography. We next placed the fossils within the modern *Podocarpus* phylogeny, following Adaimé *et al*. (2024a), and reconstructed ancestral states for temperature tolerance on the tip-calibrated tree, using environmental values derived from modern *Podocarpus* species distributions and fossil paleogeographic locations. This led to a significantly different result from the ancestral state temperature reconstructions based solely on extant species.

By integrating deep learning, morphological analyses, and phylogenetic inference, this study provides novel insights into how *Podocarpus* pollen morphology has evolved in response to changing environments over time. The abstract morphological features extracted by CNNs capture key morphological differences among species and allow us to test for pollen traits that may have been selected for under different environmental conditions and hypothesize about environmental adaptations in the pollen of *Podocarpus*. Extending these methods to fossil pollen specimens offers a framework for inferring the environmental tolerances of extinct deep-time morphotypes and suggests potential scenarios for how temperature and solar radiation tolerances evolved within the *Podocarpus* phylogeny.

## Data sources and methods

### Modern and fossil pollen specimens

We used pollen material isolated from 36 modern *Podocarpus* species, obtained as prepared slides from the Smithsonian Tropical Research Institute (STRI), Utrecht University, and the Florida Museum of Natural History. The species are distributed across Australasia/Indomalaya, the Neotropics, Afrotropics, and the Asian Palearctic (Fig. 1) (Table S1). Species names were reviewed and amended in accordance with the World Checklist of Vascular Plants (WCVP) database, accessed through Kew Plants of the World Online (2024). Each modern species was represented by a single slide, which was derived from a single herbarium specimen. Fossil *Podocarpidites* specimens were retrieved from Late Cretaceous and Neogene samples from different localities in the Neotropics, namely Colombia, Brazil, Peru, and Panama. Fossil specimens were from STRI collections and have been described in previous studies (Jaramillo *et al*., 2014; Martínez *et al*., 2020; Carvalho *et al*., 2021) (Table S2).

### Specimen imaging

A total of 375 modern pollen specimens from the 36 extant species and 31 fossil specimens (Table S2) were imaged at 630× magnification (63×/1.4 NA oil objective, 0.08 µm per pixel resolution) using a Zeiss LSM 880 equipped with Airyscan confocal superresolution (Sivaguru *et al*., 2018; Romero *et al*., 2020). Images were acquired as a series of axial focal planes captured at 0.19 µm increments between each plane. We used three distinct imaging modalities for our classification models, following the approach outlined in (Adaimé *et al*., 2024a): image cross-sections, maximum intensity projections (MIPs), and image patches. For the modern reference specimens, we created cross-sectional images by dividing the image stack into smaller segments. Each substack consisted of 15 axial planes (total depth of 2.85 µm), offset by 10 planes (1.90 µm) between cross-sections. For fossil specimens, which were more compressed, we used individual planes from the image stack as cross-sections. Maximum Intensity Projections (MIPs) were generated from the full image stack to represent the overall external structure and shape of each pollen grain. Contrast-limited adaptive histogram equalization (CLAHE) (Pizer *et al*., 1990) was applied to standardize pixel intensity values. Finally, each MIP was divided into 10-13 square patches covering ∼10% of the total pollen grain area. These patches allowed the model to focus on the textural patterning of our pollen grains.

### Reference phylogeny

Our reference phylogeny is based on the published fossil-calibrated phylogeny of 20 extant Podocarpaceae genera (Khan *et al*., 2023). The tree was generated using Bayesian inference under the fossilized birth-death (FBD) model and constructed from six genetic markers: rbcL, matK, trnL-trnF, NEEDLY, PHYP, and ITS, using the GTR+I+G nucleotide substitution model. Most clades in the phylogeny are well-supported and the phylogeny was calibrated using 20 macrofossils dated from the Permian to the Late Paleocene; 16 of these were Podocarpaceae (Khan *et al*., 2023). We used a second time-calibrated phylogeny (Leslie *et al*., 2018a) and the AddTip function of the TreeTools package in R (Smith *et al*., 2024) to add *P. glaucus* into our phylogeny as sister to *P. spinulosus*. The divergence time between *P. glaucus* and *P. spinulosus* in the new composite tree was consistent with the divergence time in Leslie *et al*. (2018). Thirty-two of the 36 modern *Podocarpus* species were included in the composite phylogeny.

### Environmental data

We inferred the environmental tolerances of the 32 modern *Podocarpus* species represented in the composite phylogeny using 14,120 occurrences in the Global Biodiversity Information Facility (GBIF) database (GBIF, 2024) (Fig. 1). To minimize potential reporting inaccuracies, we filtered the occurrence data using the geographic centroids of each species’ distribution. Occurrences located more than 7000 km from their respective centroids were considered outliers and removed. For each remaining specimen, we used the specimen’s geographic coordinates to extract mean annual temperature, annual precipitation, altitude, and solar radiation values from 30s-resolution rasters (GeoTIFF files) from the WorldClim 2 database (Fick & Hijmans, 2017) (Fig. 2).

The inferred paleotemperature data for the *Podocarpidites* fossils were retrieved from the PALEOMAP Project based on their fossil age and paleocoordinates (Scotese, 2021) (Fig. 3). We used the R package “rgplates” (Kocsis & Raja, 2019; Kocsis *et al*., 2021) to calculate paleocoordinates from present-day locations and plate tectonic models (Table S2).

**Figure 3.**
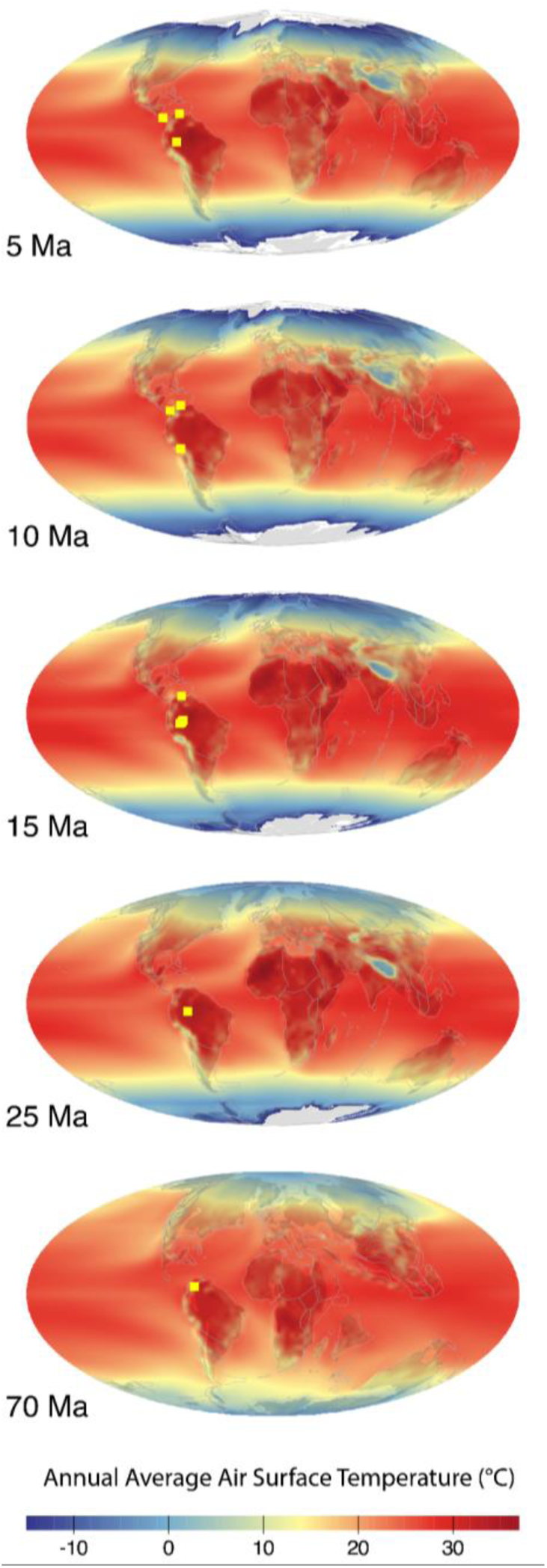
Reconstructed paleotemperature maps for five geological time periods (5 Ma, 10 Ma, 15 Ma, 25 Ma, and 70 Ma), showing the annual average air surface temperature. Yellow squares indicate the locations of fossil *Podocarpidites* specimens included in our analysis. Temperature reconstructions were generated using PALEOMAP (Kocsis *et al*., 2021; Scotese, 2021) representing the environmental conditions in which the fossils were deposited.

### Classification model training, evaluation, and feature extraction

We trained three separate convolutional neural networks (CNNs) to classify all 36 modern *Podocarpus* species across the three image modalities as described in the section on specimen imaging (Table S1). The first CNN was trained with the cross-sectional images; the second CNN was trained with MIP images; and the third CNN was trained with the image patches. Each CNN was built on a ResNeXt-101 architecture (Xie *et al*., 2017), pretrained on the ImageNet-1K dataset (Deng *et al*., 2009) and fine-tuned with our *Podocarpus* image dataset. The classification models produced *K*-dimensional probability vectors representing the likelihood of each specimen belonging to one of the *K* taxa. The models also produced 2048-dimensional feature vectors from their penultimate layers, capturing more nuanced representations of pollen morphology across each modality.

Training and validation data were split randomly, with 70% or the specimens used for training and 30% for validation. Common data augmentation techniques were applied, including random vertical and horizontal flips (P = 0.5) and random rotations within the range of [-90°, +90°] (P = 0.5). All images were automatically resized to 224×224×3 pixel resolution using bilinear interpolation. We trained the CNNs with a batch size of 10, using a cross-entropy loss function, and stochastic gradient descent optimization with momentum of 0.9 and a learning rate of 0.015. A step learning rate scheduler was used, decreasing the learning rate by half every two epochs. We trained the models for 30 epochs, and fused the predictions from the three CNNs to produce a final classification score for each specimen, assuming conditional independence between the modalities the class label (Chen *et al*., 2022; Adaimé *et al*., 2024a) and a uniform prior with respect to the class labels.

We first evaluated the classification performance of each trained CNN to ensure the models effectively recognized species-level morphological features. We then extracted the 2048-dimensional abstract features from the penultimate layer of each of our three CNNs, corresponding to the internal structure, shape, and textural patterning of the pollen grains, respectively. For the cross-sectional CNN and patch CNN, the feature values were averaged elementwise across all cross-sectional images or patches for each pollen grain. These feature vectors were then concatenated horizontally into a final 6144-dimensional feature vector for each grain, capturing fine-scale morphological information for each specimen. The concatenated feature vectors were averaged by species, reducing specimen-level differences and allowing for a more generalized representation of each *Podocarpus* species.

We used principal component analysis (PCA) on the raw CNN feature matrix to reduce dimensionality and identify the key axes of variation in pollen morphology. We used the elbow method to identify the optimal number of principal components to retain, using the “findPC” function implemented in the R package of the same name (Zhuang *et al*., 2022). Specifically, we identified the point where the second derivative of the explained variance curve approached zero, indicating a slower rate of change in explained variance (the first derivative).

### Morphospace reconstructions

We constructed the phylomorphospace of modern *Podocarpus* species along PC1 and PC2, derived from the PCA of the CNN-derived features. We generated the 95% confidence ellipse, calculated using the covariance matrix of modern species scores. This represented the most likely phenotypic range of modern *Podocarpus* pollen. Fossil features were then projected onto this PCA morphospace to compare their morphology against the range of modern taxa.

To assess how much morphological variation was constrained by the phylogenetic history of *Podocarpus*, we calculated Pagel’s λ for each retained PC axis (Pagel, 1999) using the “phylosig” function of the R package “phytools” (Revell, 2024). The significance of λ was determined using a likelihood ratio test, comparing the fit of the observed λ to a null model where λ = 0 (no phylogenetic signal).

### Relationships between pollen morphology and environmental variables

We next applied both standard linear regression and phylogenetic generalized least squares (PGLS) (Revell, 2010) to correlate PC1, PC2, and PC3 with mean annual temperature, annual precipitation, altitude, and solar radiation. For the PGLS analysis, we used the “pgls” function of the R package “caper” (Orme *et al*., 2011). We applied a Bonferroni correction (Abdi, 2007) to adjust for multiple comparisons and reduce the risk of false positives. Only relationships with corrected p-values < 0.05 were considered statistically significant.

To test for relationships between pollen length and temperature tolerance, we performed PGLS and linear regression analyses between the two variables. Specifically, we measured the total length of each pollen grain using Feret’s diameter, calculated the median grain length for each species, and then used these median values to assess the strength and significance of the relationship between grain dimensions and temperature tolerance.

Likewise, to test the hypothesis that *Podocarpus* pollen wall thickness will increase with the amount of solar radiation in the environment, we performed the same analyses to compare solar radiation values and the pixel intensity of individual pollen grains. We used pixel intensity as a proxy for corpus wall thickness. Because pollen exine is autofluorescent, all other conditions being equal, higher pixel intensity values indicate a thicker and more robust pollen wall (Sivaguru *et al*., 2018; Adaimé *et al*., 2024b). We computed the median pixel intensity for each specimen and averaged these values per species to estimate the overall pollen wall thickness for each species.

We used the PC1 scores for each fossil specimen, projected onto the morphospace defined by the 32 modern species, to calculate their estimated temperature value from the PLGS regression, based on the best-fit line between the modern *Podocarpus* PC1 scores and mean annual temperature. The fossil PC1-inferred temperature values were then compared to the reconstructed temperatures for each fossil derived from PALEOMAP (Kocsis *et al*., 2021; Scotese, 2021). We used log-log linear regression to address potential nonlinearity between the two variables and stabilize the variance and Pearson’s correlation coefficient to determine the strength and significance of the relationship. We visualized the placement of the fossil specimens on the PGLS regression by plotting their PC1 scores relative to their PALEOMAP temperatures (Kocsis *et al*., 2021; Scotese, 2021).

### Phylogenetic placement of fossil *Podocarpidites* specimens

#### Multilayer Perceptron (MLP)

Following the approach described in Adaimé *et al*. (2024a), we used a multilayer perceptron (MLP) to transform the morphological features into phylogenetically-informed features. The MLP took as input the 6144-dimensional feature vector, obtained by concatenating the 2048-dimensional vectors from each of the three CNNs trained on different pollen image modalities. The MLP reduced the difference between the pairwise distances in the feature space and the cophenetic distances derived from the time-calibrated reference phylogeny. The MLP was trained with a loss function that minimized the mean squared error (MSE) between the predicted pairwise distances from the output feature vectors and the evolutionary distances derived from the phylogeny.

The MLP consisted of four fully connected layers, with the number of neurons gradually decreasing from the original 6144 in the input layer to 256 in the final output layer. Dropout was applied after the second and third layers, with a dropout rate of 0.5 to prevent overfitting. A rectified linear unit (ReLU) activation function was applied after each layer to introduce non-linearity (Nair & Hinton, 2010). The model was trained using the AdamW optimizer with a learning rate of 0.0001 and weight decay of 10^-4^. After training over 10,000 iterations, the MLP produced a 256-dimensional embedding for each species, representing the phylogenetically-informed morphological features to be used in phylogenetic placement.

#### Tree construction

We first forward-passed the fossil pollen images through each of the three trained CNNs, concatenating the resulting feature vectors across the three modalities for each fossil specimen. These concatenated vectors were then input into the MLP, which had been trained to transform the features according to a phylogenetically informed distance function. We extracted the resulting 256-dimensional output feature vectors and used them together with the species-level MLP feature vectors of modern species as the final dataset for inferring the phylogeny. We used these embedding features to construct the phylogenetic tree under Bayesian Inference, assuming a Brownian motion (BM) model for character evolution. Analyses were conducted in RevBayes (v.1.2.4) (Höhna *et al*., 2016), with Markov Chain Monte Carlo (MCMC) simulations run for 1 million generations, split into two independent runs, each with four chains. Trees were sampled every 1000 generations, and burn-in was determined through visual inspection of convergence using Tracer (v.1.7.1) (Rambaut *et al*., 2018). The maximum clade credibility (MCC) tree was generated using TreeAnnotator (v.2.4.2) and selected as the final phylogeny, with branches considered significantly supported if their posterior probabilities were ≥ 0.95. We visualized and edited the final trees using the function “plotTree” of the R package “RevGadgets” (v.1.1.1) (Tribble *et al*., 2022).

Diversification and turnover rates were drawn from uniform distributions ranging from 0.5 to 1.5 and 0.5 to 2, respectively, with speciation defined as the sum of these rates. The fossilization rate was drawn from a uniform distribution ranging from 0.05 to 0.1. The extinction rate was set equal to the speciation rate, and the sampling fraction was fixed at 0.2759, representing the fraction of *Podocarpus* species included in our dataset. The root age was drawn from a uniform distribution ranging from 67.2 Ma, the maximum age of our oldest fossil type, to 82 Ma, based on the maximum root age inferred for *Podocarpus* inferred by Quiroga *et al*. (2016).

### Ancestral state reconstructions of environmental distributions

We reconstructed the ancestral states for mean annual temperature, annual precipitation, altitude, and solar radiation across the modern *Podocarpus* phylogeny, based on the WorldClim averages for each species (Fick & Hijmans, 2017) (Fig. 2). To model the evolution of these environmental tolerances, we fit three models of evolution to our data: BM, early burst (EB), and Ornstein-Uhlenbeck (OU) with a moderate α (representing the rate of attraction towards an optimum trait value) using the “fitContinuous” function from the R package “geiger” (Pennell *et al*., 2014; Harmon *et al*., 2023). We selected the best evolutionary model based on the lowest Akaike information (AIC) criterion and highest likelihood values, and then used the best fitting model to perform ancestral state reconstruction using the “reconstruct” function from the R package “ape” (Paradis *et al*., 2002; Paradis & Schliep, 2019). This provided us ancestral state reconstructions based solely on extant distributions. We repeated the analysis using our tip-calibrated tree, which incorporated our fossil pollen specimens and their ages. This allowed us to directly incorporate fossil data into our ancestral state reconstruction, including their inferred paleotemperature tolerances, and compare the results to those based solely on extant species.

We estimated phylogenetic signal in environmental tolerances for the four environmental variables by calculating Pagel’s λ (Pagel, 1999) using the “phylosig” function of the R package “phytools” (Revell, 2024). We tested for significance using a likelihood ratio test, comparing the observed λ to a null model where λ = 0. P-values were obtained from 1,000 randomizations.

## Results

### Classification accuracy

The CNNs trained on cross-sectional images, MIPs, and patches, achieved classification accuracies of 89.34%, 75.41%, and 68.03%, respectively (Fig. S2). Combining the predictions of all three models resulted in the highest overall accuracy (90.16%) (Fig. S2). We averaged the feature values across species and concatenated the feature vectors from all three CNNs for use in subsequent analyses. This is the “untransformed feature matrix” referenced throughout the results. All 36 extant species were used in training the CNNs, but only the 32 species represented in the reference phylogeny were included in the feature matrix and subsequent analyses.

#### Morphospace analyses

The phylomorphospace reconstructions of the untransformed CNN features show a clear separation between the two subgenera of *Podocarpus* (*P.* subg. *Podocarpus* and *P.* subg. *Foliolatus*) (Fig. 4), particularly along PC2. Notably, PC2 showed moderate but significant phylogenetic signal (Pagel’s λ = 0.46, p = 0.0002). In contrast, PC1 (λ = 0.13, p = 0.41) and PC3 (λ ≈ 0, p = 1) exhibited little to no phylogenetic signal. The projection of fossil features onto the phylomorphospace, defined by the PCA factor map of modern *Podocarpus* pollen, indicates that all fossils lie within the 95% confidence ellipse of the expected range of morphological variability in modern *Podocarpus* pollen (Fig. 4B).

**Figure 4.**
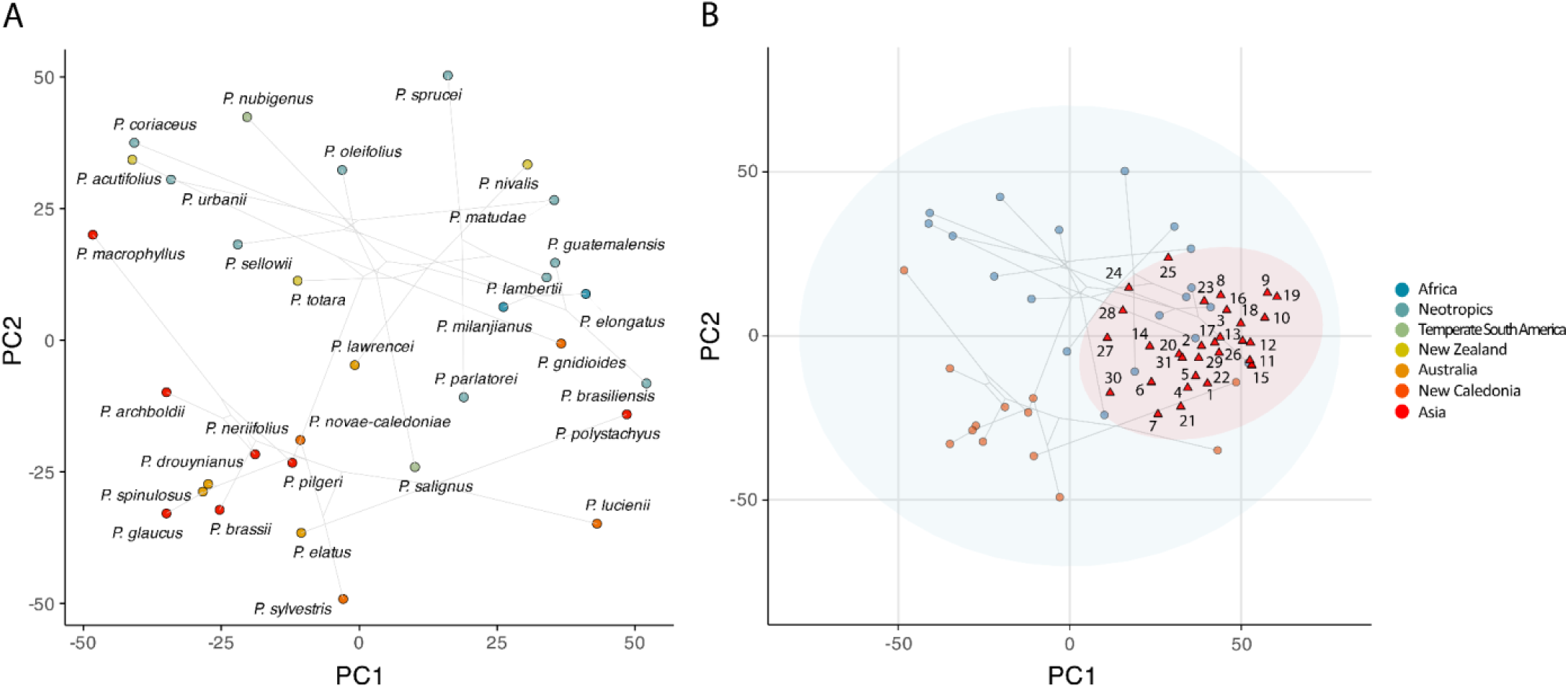
(A) Phylomorphospace visualization of PCA1 and PCA2 scores obtained from the PCA conducted on the CNN feature matrix. Each point represents a *Podocarpus* species, with colors corresponding to their biogeographic regions (see Figure 1). Lines connecting the points represent the phylogenetic relationships between species (based on Khan *et al*., 2023), indicating the species’ evolutionary trajectories across the morphospace. (B) Fossil specimens projected onto the phylomorphospace of modern *Podocarpus* taxa. Red triangles represent the fossil specimens. Nodes are colored by subgenus, with species from subgenus *Podocarpus* colored in blue and those of subgenus *Foliolatus* colored in orange. The blue ellipse represents the 95% confidence interval for the predicted morphospace range of modern species, while the red ellipse represents the expected morphospace range occupied by the fossils.

### Relationships between pollen morphology and environmental variables

We retained the first three principal components of the untransformed feature matrix of the 32 species represented in the reference phylogeny based on the results of the elbow method (Fig. S3). These three components accounted for 35.97% of the total variability in the dataset (PC1:15.3%, PC2: 12.15%, PC3: 8.49%).

PC1 and mean annual temperature showed a significant correlation in PGLS (adjusted R^2^ = 0.3056, p = 0.0006134, adjusted p = 0.00736), although the linear regression model for this relationship was not significant (Fig. 5A). Further analysis revealed that pollen grain size and mean annual temperature were significantly correlated in both PGLS (adjusted R^2^ = 0.1956, p = 0.008357) and linear regression (adjusted R^2^ = 0.153, p = 0.01865) (Fig. 5B).

**Figure 5.**
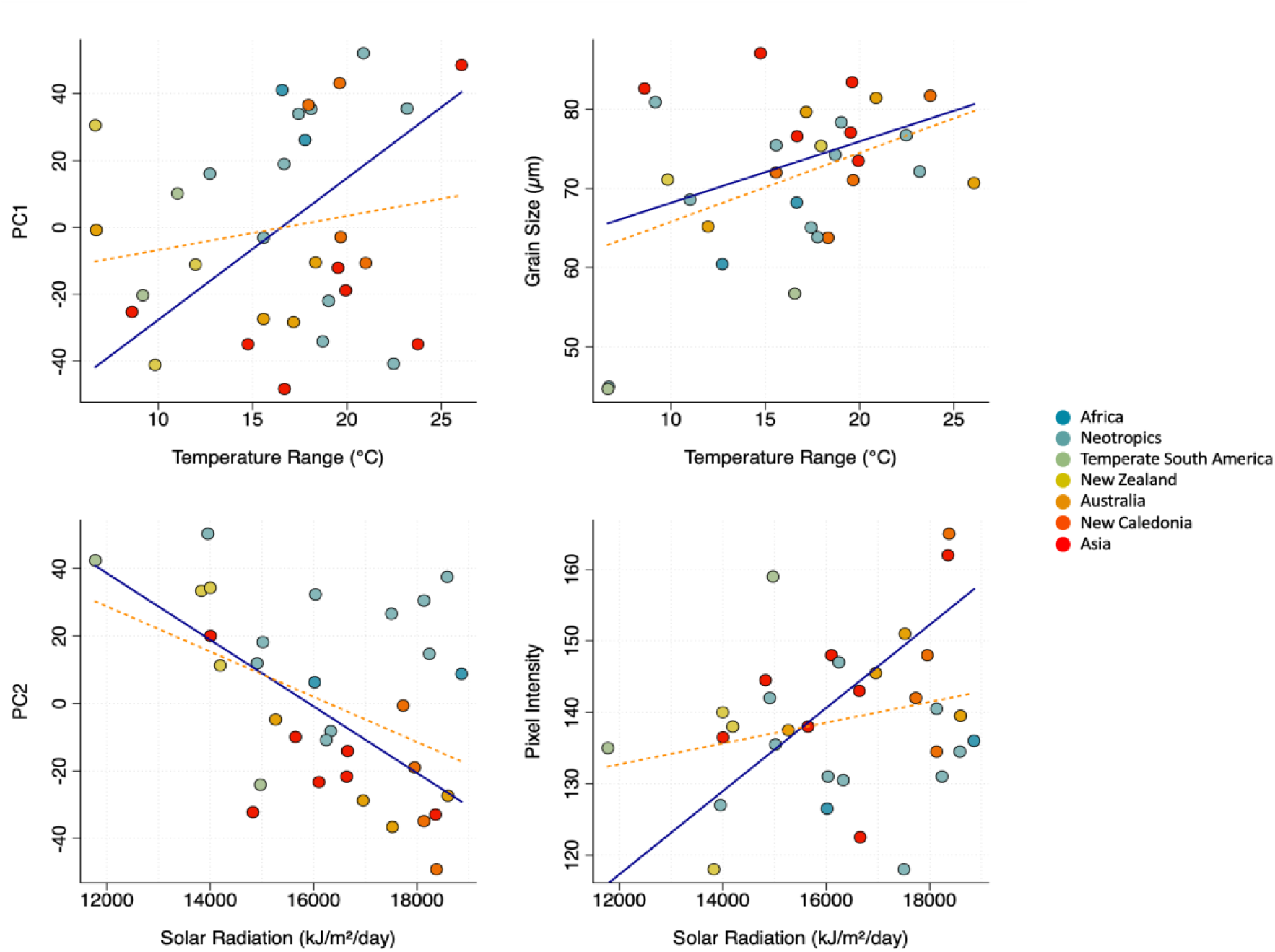
Phylogenetic Generalized Least Squares (PGLS) and standard linear regression models (LM), illustrating the relationships between the two selected environmental variables (i.e. temperature range and solar radiation) and morphological variation in *Podocarpus* pollen. Solid blue lines represent the PGLS regression fits, while dotted orange lines represent the LM fits. The top-left panel shows a significant positive relationship between PC1 and mean annual temperature in the PGLS analysis, indicating that species from warmer regions exhibit higher PC1 scores. The top-right panel shows a positive relationship between total grain size and temperature, with both PGLS and LM analyses identifying significant relationships. The bottom-left panel shows a significant negative correlation between PC2 and solar radiation in the PGLS analysis, with significant results in both PGLS and LM, suggesting that species in high solar radiation environments tend to exhibit lower PC2 scores. The bottom-right panel reveals a significant positive relationship between pixel intensity and solar radiation in the PGLS analysis, suggesting that species exposed to higher solar radiation exposure tend to have pollen grains with thicker corpus wall. Species are color-coded by biogeographic region. PGLS significance was determined after applying a Bonferroni correction.

PC2 and solar radiation showed significant correlations in both PGLS (adjusted R^2^ = 0.2424, p = 0.002472, adjusted p = 0.0297) and linear regression (adjusted R^2^ = 0.1685, p = 0.01131) (Fig. 5C). The relationship between solar radiation and corpus wall thickness, inferred from median pixel intensity, was significant only in PGLS (adjusted R^2^ = 0.2761, p = 0.001189) (Fig. 5D), suggesting that thicker pollen walls are associated with higher solar radiation levels across *Podocarpus* species, after correcting for phylogenetic relatedness.

When we inferred the temperature tolerances of the fossil *Podocarpidites* specimens using their projected PC1 scores and the PGLS regression model (Fig. 6), the analysis revealed two distinct clusters: one of the southern Peruvian Altiplano (12 Ma) specimens with lower PC1 scores, associated with cooler temperatures (∼15 °C), and a second cluster of specimens from lowland Brazil, Panama, Venezuela, and Colombia (3.6-16.5 Ma) with higher PC1 scores, corresponding to warmer temperatures (∼20-31 °C). When we compared the predicted temperatures from the PGLS model to PALEOMAP reconstructed temperatures (Kocsis *et al*., 2021; Scotese, 2021) using log-log regression, we found a moderate but significant positive correlation (r = 0.3809, p = 0.0345) (Fig. S4).

**Figure 6.**
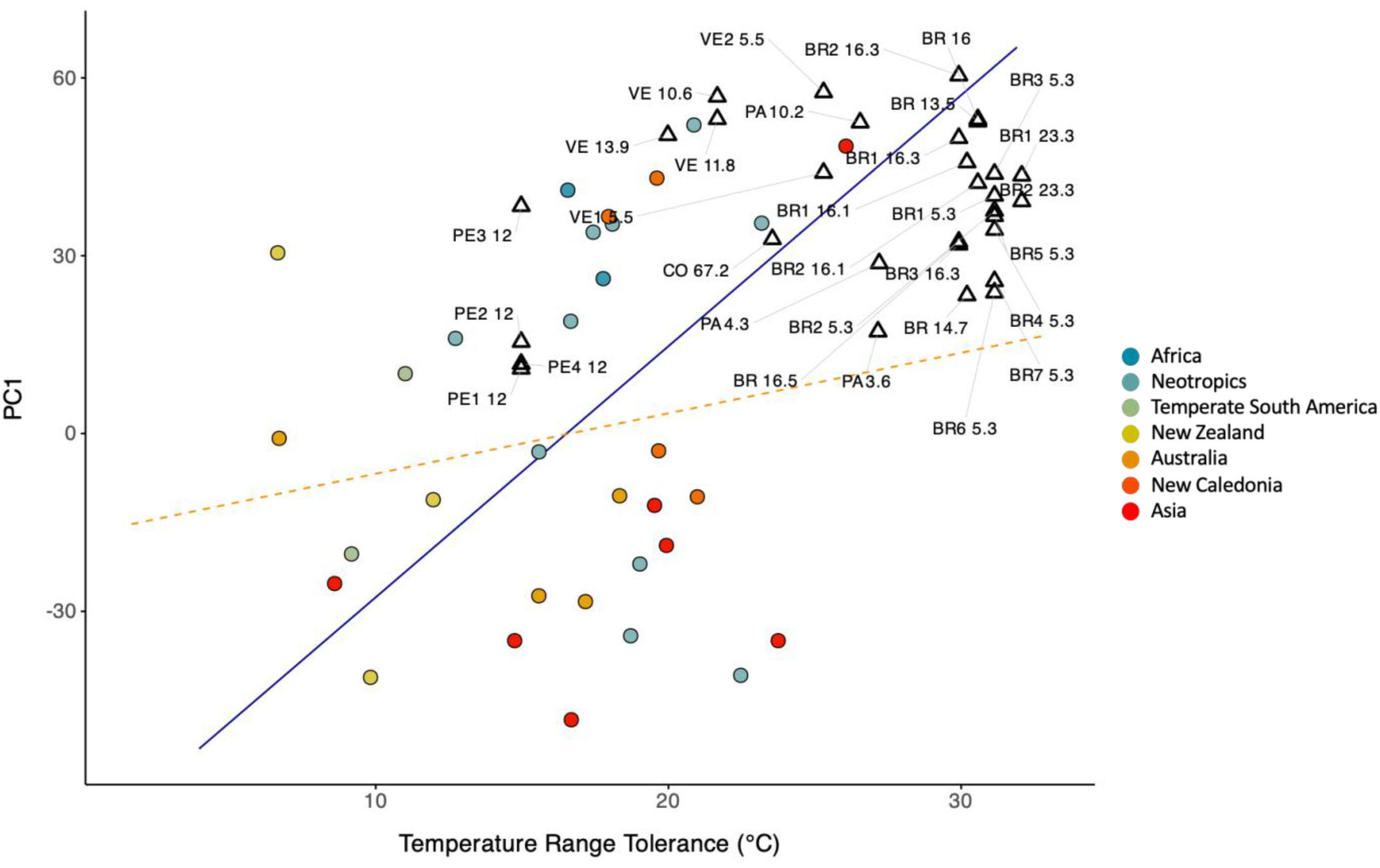
Projection of fossil data onto the linear regression- and PGLS-derived trendline between PC1 and temperature based on modern *Podocarpus* species (color coded by geographic region). Two distinct clusters emerge among the fossil data: cold-tolerant Peruvian (PE) fossils, which exhibit lower PC1 scores and are associated with lower reconstructed paleotemperatures of 15 °C (Kocsis *et al*., 2021; Scotese, 2021), and a cluster comprised of the remaining fossils from Brazil (BR), Panama (PA), Venezuela (VE), and Colombia (CO), displaying higher PC1 scores and associated with warmer reconstructed paleotemperatures, ranging from 20 to 31 °C. The dashed line represents the PGLS trendline derived from the modern data.

### Phylogenetic placement of fossil *Podocarpidites*

Eighteen of the fossil *Podocarpidites* specimens (dating from 16.3 to 3.6 Ma) were placed within the main Neotropical clade (P > 0.95) (Fig. S5). Three out of the four fossil specimens from Altiplano southern Peru (12 Ma) were placed as sister to the Andean Chilean species *P. salignus* (P = 1). This placement altered the original tree topology, notably making the two Chilean species *P. salignus* and *P. nubigenus* sister taxa, and outgroup to the other Neotropical and African species. The Late Cretaceous Colombian fossil (67.2 Ma) and five Miocene samples from Brazil and Venezuela (dating from 16.5 to 5.3 Ma) were placed as sister to the entire subgenus *P. Podocarpus* (P = 0.9973).

Three out of the 31 fossil specimens were placed as either sister or within the Australasian subgenus *P.* subg. *Foliolatus*. Specifically, two Middle and Late Miocene specimens (11.8 and 5.3 Ma) from Venezuela and Brazil were placed as sister to subg. *Foliolatus* (P = 1 and P = 0.8564, respectively), and a Late Miocene morphotype from Brazil (5.3 Ma) was placed with the lineage of *P. elatus* and *P. polystachyus* (P = 1). We consider these placements questionable outliers given the accepted biogeographic history of the genus (Quiroga *et al*., 2016).

### Ancestral states for environmental tolerances

The Ornstein-Uhlenbeck (OU) model was the best-fitting model for ancestral state reconstructions, based on living species, across all four environmental variables: mean annual temperature, annual precipitation, altitude, and solar radiation. OU model reconstructions suggest that early *Podocarpus* lineages may have been adapted to cooler temperatures and moderate precipitation, before diversifying into a broader climatic niche (Figs. 2A and 2B). Additionally, reconstructed ancestral states indicate adaptations to lower elevations and higher levels of solar radiation (Fig. 2C).

However, ancestral state reconstructions that integrated fossil placements and PALEOMAP reconstructed temperatures changed the reconstructed node values for the evolution of temperature tolerance across the genus (Fig. 7). In these reconstructions, *Podocarpus* likely originated as warm-tolerant, with later adaptations for cooler climates emerging independently in specific lineages, including the New Zealand and Andean lineages. The monophyletic group encompassing the South American and African lineages likely shared a warm-tolerant common ancestor (Fig. 7).

**Figure 7.**
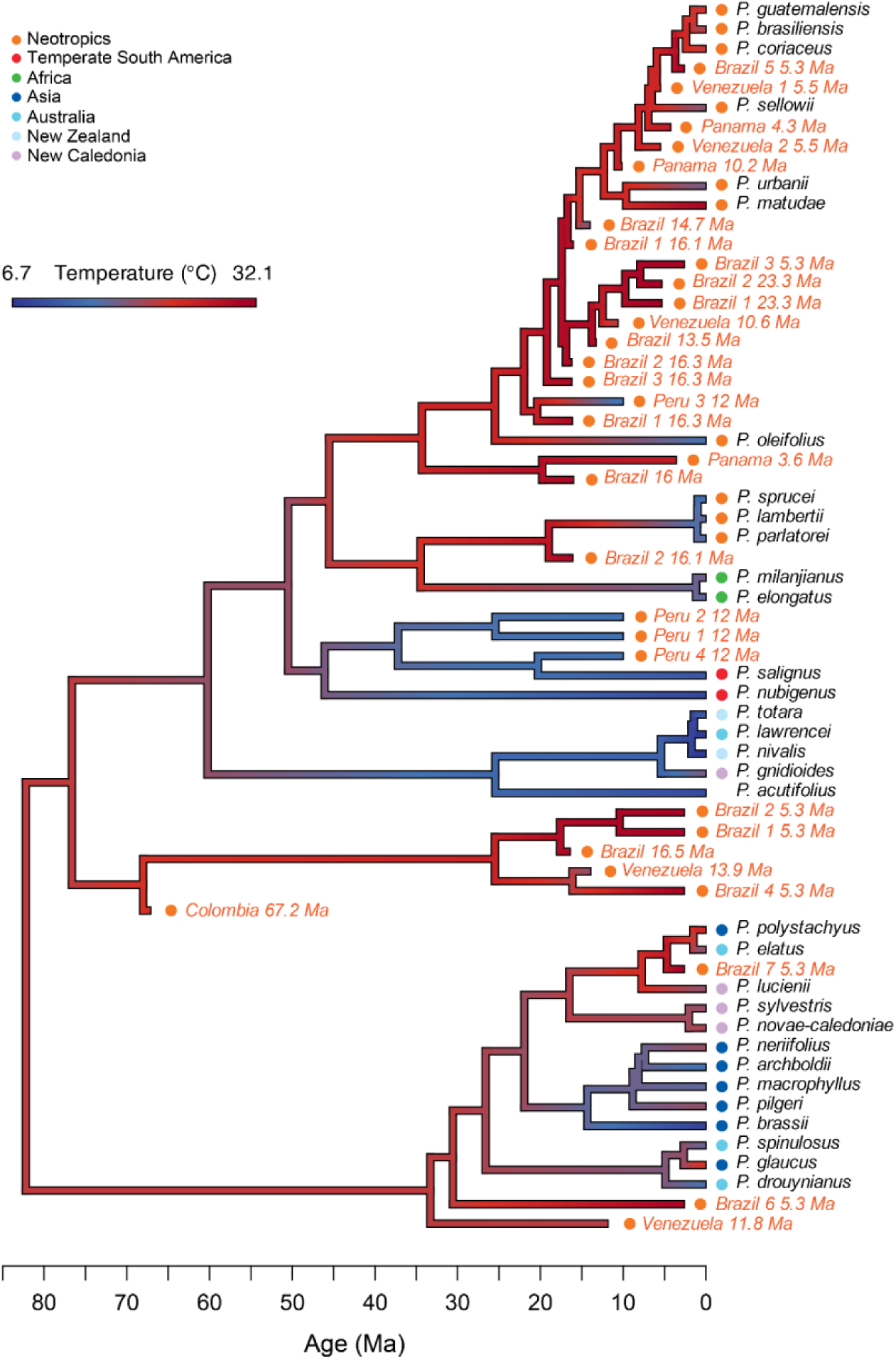
Ancestral state reconstruction of temperature using both the modern and fossil data, assuming the BM model of evolution. Branch colors represent reconstructed temperatures, ranging from cooler (blue) to warmer (red). Fossil placements are based on the Bayesian inference tree reconstructed using the approach in (Adaimé *et al*., 2024a). The inclusion of fossils supports a warm-tolerant origin for *Podocarpus* subg. *Podocarpus*, with later adaptations of specific lineages (e.g., the New Zealand and Andean clades) to cooler climates due to environmental pressures.

Pagel’s λ indicates a strong phylogenetic signal for mean annual temperature (λ = 0.73, p = 0.0084), suggesting that temperature tolerance is phylogenetically conserved across *Podocarpus*. Solar radiation also showed a high phylogenetic signal (λ = 0.73), though this result was not statistically significant (p = 0.084). In contrast, annual precipitation (λ = 0.24, p = 0.27) and altitude (λ ≈ 0, p = 1) showed weak or no phylogenetic signal, indicating that these traits are not strongly influenced by phylogenetic relatedness.

## Discussion

By integrating statistical approaches with CNN-derived features, we identified the environmental factors that shape pollen morphology in *Podocarpus* and applied these insights to infer past environmental tolerances of extinct pollen types. This approach demonstrates how pollen morphologies link with environmental variability within an evolutionary framework. *Podocarpus* pollen morphology is not strictly constrained by phylogeny, but also demonstrates adaptations to temperature and solar radiation.

Pollen morphology and temperature tolerance have not been widely examined in an evolutionary framework. However, in cross-species analyses, Ejsmond *et al*. (2011, 2015) and McCulloch *et al*., (2022) found that species with larger pollen grains are often associated with warmer climates, and suggest that this may reflect adaptations for water conservation and extended viability through reduced surface-to-volume ratios. Within a species, temperature can enhance pollen hydration and size, as in *Pinus sylvestris* (Varis *et al*., 2011). The significant correlation between the primary axis of variation in the CNN-learned pollen classification features (PC1), pollen size, and temperature suggests that our study also identified an analogous adaptive morphological response to temperature, even after accounting for phylogenetic relatedness (Figs. 5A and 5B). Our pollen images were standardized for size, so the CNN features we produced are size independent. However, PC1 may be capturing differences in texture and shape that correlate with size and may be in response to temperature fluctuations and desiccation pressure.

We found a similar significant association between pixel intensity and solar radiation (Fig. 5D). Pixel intensity serves as a proxy for corpus wall thickness under standardized imaging parameters, given that the exine, which is naturally autofluorescent, shows stronger fluorescence with increased thickness (Sivaguru *et al*., 2018; Adaimé *et al*., 2024b). This suggests that thicker pollen walls may have been selected for in environments with higher solar radiation. Similar to the findings by Yeloff *et al*. (2008) where thicker pollen walls were observed under higher UV-B conditions, this adaptation likely serves to protect the cells that form the male gametophyte and reduce DNA damage to male gametes (Ariizumi & Toriyama, 2011).

However, determining the cause behind these correlations is not straightforward. No analogous studies provide a comparative framework for investigating similar pollen adaptations over evolutionary timescales, so we are unable to directly link our observed morphological differences to long-term adaptive processes. Secondly, the pollen grains in our analysis are from a single individual per species and may not fully capture the full range of intraspecific variability present within *Podocarpus*. Because our samples are limited in this way, we also cannot rule out alternative explanations for the observed correlations between environment and pollen morphology, such as phenotypic plasticity (Ejsmond *et al*., 2011).

When we projected the CNN features of the fossil specimens onto the PCA morphospace defined by modern *Podocarpus* species, using their PC1 scores and reconstructed temperature values, the fossil samples broadly followed the temperature-PC1 relationship defined by linear regression and PGLS models, with lower PC1 scores corresponding to cooler temperatures and higher scores associated with warmer temperatures. Samples from the Altiplano in Southern Peru (12 Ma) were taken in sediments that accumulated when the region was at ∼2 Km of elevation in a montane forest (Martínez *et al*., 2020). They showed the lowest PC1 scores and the lowest reconstructed temperatures as would be expected compared to other fossil sites collected from the tropical lowlands (Fig. 6). When we used the modern PGLS trendline for PC1 as a predictive model to estimate fossil temperature tolerances and compared these predictions to PALEOMAP-derived temperature estimates using log-log regression analysis (to account for potential nonlinearity), there is a positive and significant correlation between predicted and reconstructed temperatures (r = 0.38, p = 0.035) (Fig. S4).

Despite the low temporal and spatial resolution of the PALEOMAP temperature reconstructions and the coarseness of the PC1-based predictions, our model captured a potential temperature signal in pollen morphology. This suggests that we may be able to approximate paleotemperatures from *Podocarpus* fossil pollen specimens. More importantly, it also shows that temperature may have been a selective factor that shaped *Podocarpus* pollen morphology over time.

Projecting all fossil features onto the modern *Podocarpus* pollen morphospace revealed that the fossil pollen specimens exhibit traits that fall within the 95% confidence ellipse of the modern morphospace as defined by PC1 and PC2 (which accounted for 27.48% morphological variation in all modern taxa). However, several specimens occupy areas of morphospace currently not occupied by modern *Podocarpus* (Fig. 4B, PC1 > 50). This may represent extinct pollen morphologies that reflect the warmer climates of the Cenozoic without modern analogues. The Neotropical fossils clustered most closely with the *Podocarpus Podocarpus* subgenus, indicating that there is some phylogenetic signal even within the untransformed CNN features. In previous research, we demonstrated that the phylogenetic signal in CNN features is weak (Adaimé *et al*., 2024a), but our morphospace analysis indicates that it is present. CNN features represent the morphological characteristics that best differentiate the modern species. Our results suggest that this is a combination of morphological differences reflecting both phylogeny and environmentally-influenced traits.

We transformed the CNN features with an MLP trained on phylogenetic distance (Adaimé *et al*., 2024a) to place the fossils in the time-calibrated *Podocarpus* phylogeny (Fig. S5). Warm-tolerant *Podocarpidites* fossils, with PALEOMAP-estimated temperature tolerances ranging from 22 °C (Middle Miocene samples from Venezuela) to 31 °C (Late Miocene samples from Brazil) (Kocsis *et al*., 2021; Scotese, 2021), were placed predominantly with the main Neotropical clade, whose extant lineages also thrive in warm environments. The Altiplano Peruvian fossils from the Middle Miocene (12 Ma), and derived from an Andean forest with PALEOMAP-estimated temperatures of 15°C similar to the empirical estimates of the same site using the paleobotanical record (Martínez *et al*., 2020) were placed as sister to *P. salignus*, suggesting close phylogenetic relatedness to cold-tolerant southern Andean lineages. The placement of fossil specimens within the *Podocarpus* phylogeny suggests a strong phylogenetic signal for temperature tolerance in the genus, with closely related species sharing similar tolerances. This is notable given that phylogenetic placement used different morphological features (MLP-transformed) than our environmental trait analysis (CNN-untransformed). The results also suggest that temperature may have influenced changes in pollen morphology independently of phylogenetic constraints, supporting the idea that both evolutionary history and environmental pressure shaped pollen traits.

When we incorporated fossils into the *Podocarpus* phylogeny, our interpretation of the evolution of cool-temperature tolerance in *Podocarpus* changed. Without fossils, ancestral state reconstructions suggest that early *Podocarpus* lineages were likely adapted to moderately cool climates. However, the inclusion of fossils suggests that tolerance to warmer conditions was the ancestral trait (Fig. 7). The Late Cretaceous Colombian specimen, placed as sister to the entire subgenus *Podocarpus Podocarpus*, was found in a moderately warm environment, with reconstructed temperatures of 24 °C according to paleotemperature reconstructions. Ancestral state reconstructions incorporating both modern and fossil taxa suggest that the subgenus was likely adapted to warm climates. This adaptation explains the warm tolerance observed in the highly diverse Neotropical clade, as well as the moderate warm tolerance in the African and temperate South American lineages (Fig. 7). Conversely, cold tolerances seen in the New Zealand and Andean clades, including the southern Peruvian fossil taxa from the Middle Miocene (Fig. 7), appear to have resulted from independent shifts from warm to cold tolerance.

Warm tolerance as the ancestral trait for *Podocarpus* is supported by the estimated origination date of the genus during a particularly warm period in Earth’s history ∼58 Ma (Khan *et al*., 2023). In South America, this period is marked by a rapid increase in species diversity, with the crown ages of many tropical clades dated between 59 and 56 Ma (Jaramillo, 2023). Our results supporting the warm-tolerant origins of the *Podocarpus* subgenus clade align with Quiroga *et al*. (2016), who suggested the last common ancestor of the subgenus inhabited warm tropical South American climates, with cold-tolerant lineages adapting to cooler habitats in the early Oligocene. Ancestral state reconstructions for *Podocarpus* subg. *Foliolatus* may be less reliable due to the questionable placement of the three Neotropical fossils within this subgenus.

Our study demonstrates that abstracted features derived from convolutional neural networks, while powerful for classifying species and placing extinct pollen morphotypes in a reference phylogeny, also provide valuable insights into evolutionary adaptations through time. The integration of fossil data, phylogenetic placements, and environmental reconstructions, enhances our understanding of how environmental variability, such as temperature and solar radiation, in conjunction with phylogenetic constraints, may have shaped the evolution of this diverse genus. Deep learning enables unprecedented exploration of evolutionary adaptations in plant lineages by quantifying complex pollen morphological traits, overcoming past limitations in capturing fine-scale morphological diversity in pollen and linking these traits to environmental change.

## Data Availability

The *Podocarpus* and *Podocarpidites* images used in the study are available from Punyasena et al. (2023) and Punyasena et al. (2025).

## Acknowledgments

We thank Hongshan Wang (Florida Museum of Natural History) and Timme Donders (Utrecht University) for providing us with microscope slides of modern Podocarpus. M-EA was supported by a University of Illinois College of Liberal Arts and Sciences COVID Recovery Grant to SWP and the University of Illinois Phillips Memorial Fund for Paleobotany. Imaging costs and postdoctoral support for MAU were provided by NSF-DBI – Advances in Bioinformatics (NSF-DBI-1262561 to SWP). SK was supported in part by the University of Macau (SRG2023-00044-FST).

## Author Contributions

M-EA conducted the machine learning and statistical analyses, supervised by SK and SWP. M-EA, SK, and SWP designed the research. MAU and M-EA collected the microscopy images. CJ identified the *Podocarpidites* material used in study. M-EA and SWP wrote the paper with feedback from all authors.

