## Supplementary Information for "Pollen morphology, deep learning, phylogenetics, and the evolution of environmental adaptations in *Podocarpus*"

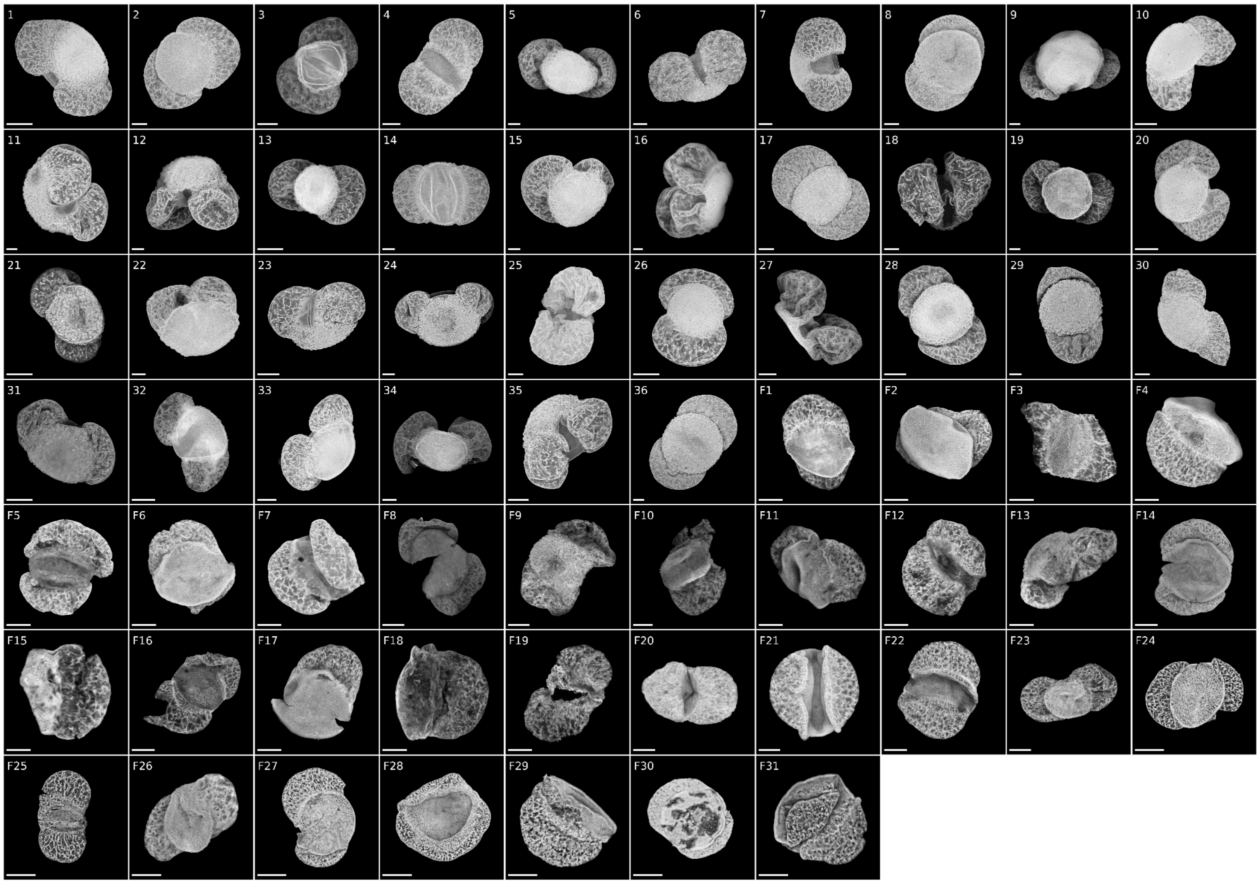

**Figure S1.** Example maximum intensity projection (MIP) images of the 36 modern species and 31 fossil specimens of *Podocarpus*. MIPs are derived from image stacks taken using optical superresolution (Zeiss LSM 880 with Airyscan, 63 × /1.4NA objective). Plate numbers and their corresponding species or specimens can be found in Tables S1 and S2 (scale bar: 10 μm).

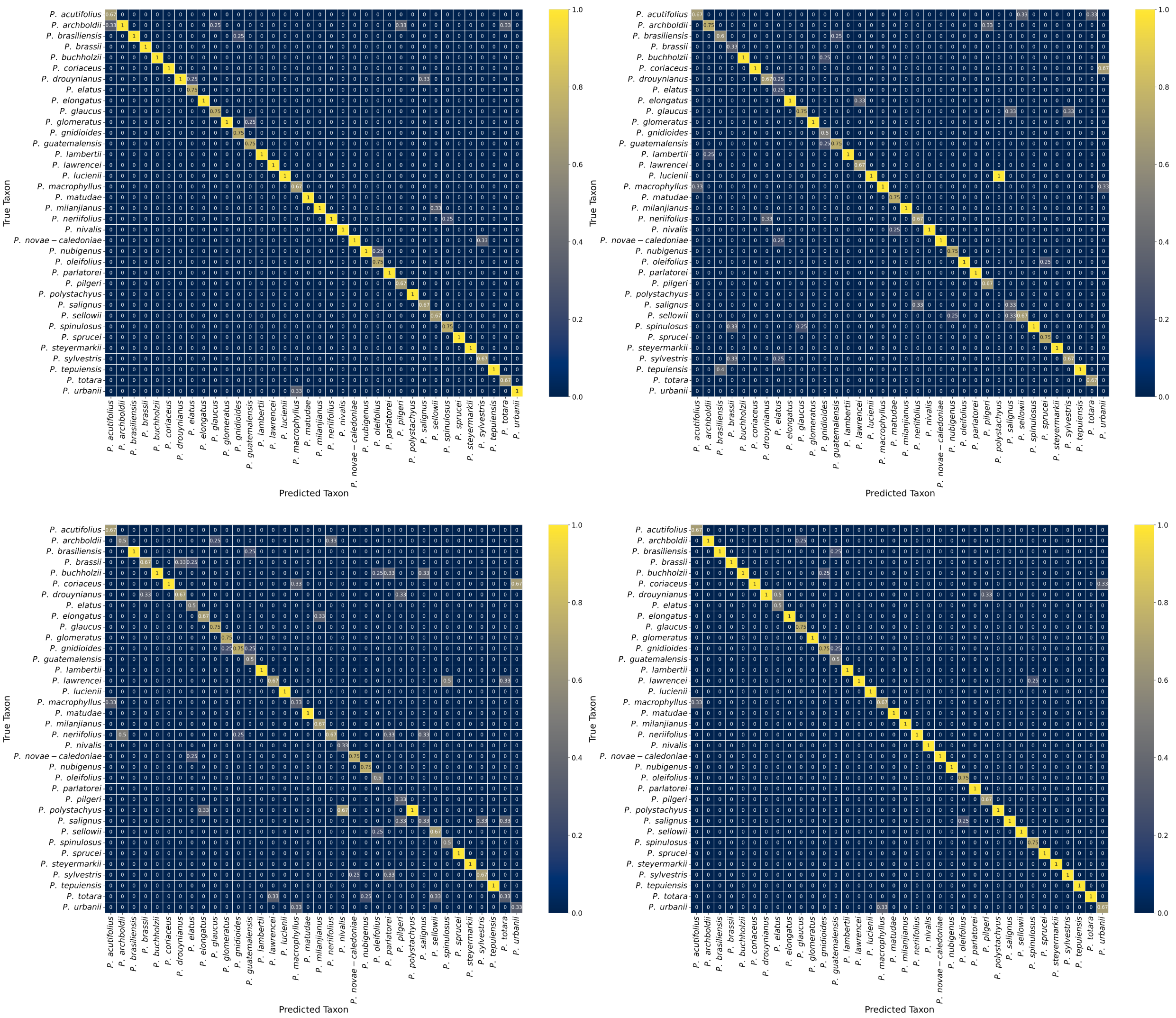


**Figure S2.** Confusion matrices illustrating the classification accuracy of the C-CNN (top left), H-CNN (top right), and P-CNN (bottom left), alongside the results of the fusion of predictions from all three models (bottom right), for the 36 modern *Podocarpus* species. Rows represent the true species labels, while columns represent the predicted species classifications by the model.

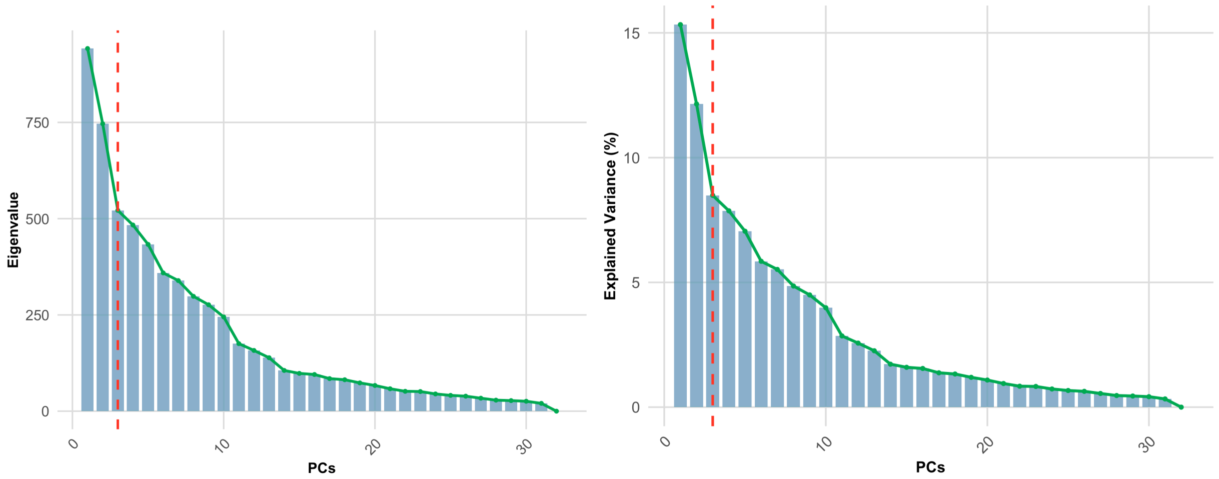


**Figure S3.** Scree plots summarizing the results of PCA. The left panel shows the eigenvalues for each PC, indicating the total amount of variance represented by each component. The right panel shows the percentage of variance explained by each PC. The dashed line marks the identified “elbow point” identified using the second derivative method, indicating the optimal number of PCs to retain.

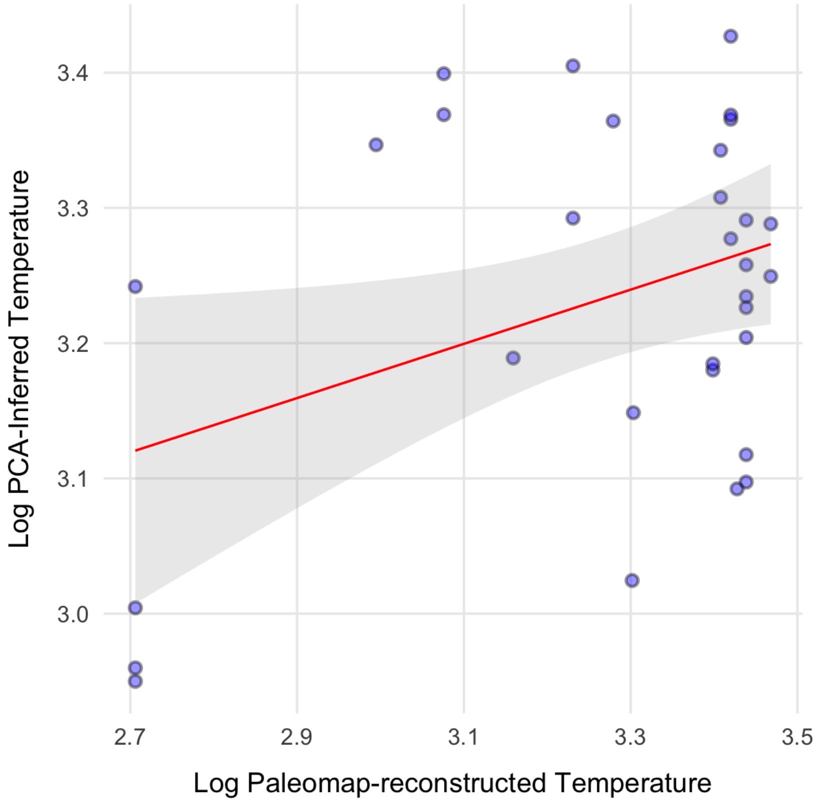


**Figure S4.** Log-log linear regression model showing the relationship between log-transformed reconstructed temperatures (obtained from PALEOMAP; Kocsis *et al.*, 2021; Scotese, 2021) and log-transformed predicted temperatures. Predicted temperatures were estimated by projecting fossil PC1 scores onto the PGLS model linking PC1 scores to mean annual temperature tolerances for modern *Podocarpus* species. The red regression line indicates a statistically significant positive correlation between the predicted and reconstructed temperatures, suggesting that the predictions based on pollen morphology align with paleotemperature reconstructions. The shaded area represents the 95% confidence interval for the regression.

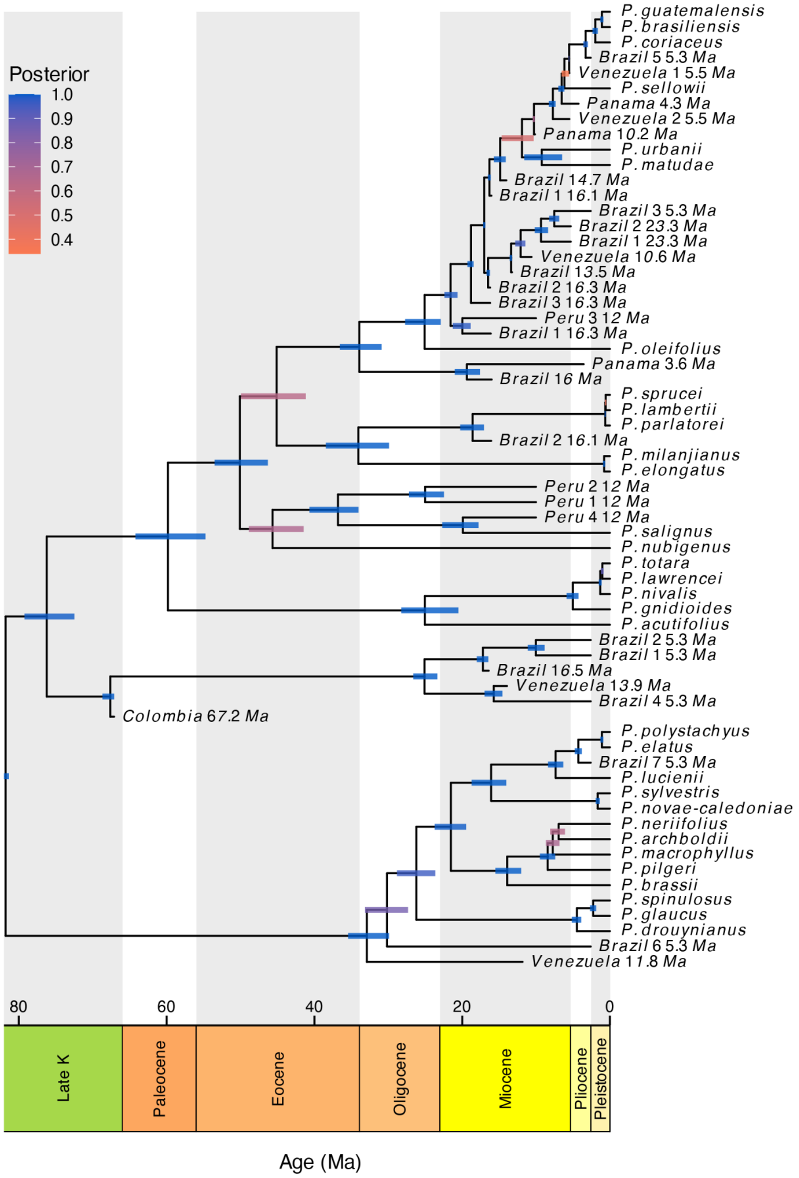


**Figure S5.** Tip-calibrated fossil tree simulated under the fossilized birth-death (FBD) process using phylogenetically informative embedding features. Posterior probability values are represented by colored bars. The tree topology closely follows that of the reference phylogeny (Khan *et al.*, 2023), with minor topological adjustments introduced due to the inclusion of fossil data and tree recomputation. Branch lengths were estimated based on our fossil ages. The resulting tree was used to conduct ancestral state reconstructions of temperature tolerance evolution in *Podocarpus*, incorporating both modern and fossil tolerance data.

**Table S1.** Modern *Podocarpus* reference specimens. Images from Punyasena et al. (2023, 2025).

| Species | No. Specimens | Geographic Range | Collection | Slide ID |
|---|---|---|---|---|
| *Podocarpus acutifolius* | 10 | Aust | STRI - Graham | 1675959 |
| *Podocarpus archboldii* | 11 | Aust | STRI - Graham | 19848 |
| *Podocarpus brasiliensis* | 14 | Neo | STRI - CTPA | 896 |
| *Podocarpus brassii* | 10 | Aust | STRI - Graham | NA |
| *Podocarpus buchholzii* | 11 | Neo | STRI-CTPA | 1389 |
| *Podocarpus coriaceus* | 10 | Neo | STRI - Graham | 845414 |
| *Podocarpus drouynianus* | 10 | Aust | STRI - Graham | 19836 |
| *Podocarpus elatus* | 11 | Aust | STRI - Graham | 19837 |
| *Podocarpus elongatus* | 10 | Afr | Utrecht | 1540 |
| *Podocarpus glaucus* | 11 | Aust/Indomalay | STRI - Graham | 19847 |
| *Podocarpus glomeratus* | 12 | Neo | STRI-CTPA | 1390 |
| *Podocarpus gnidioides* | 12 | Aust | STRI - Graham | 22388 |
| *Podocarpus guatemalensis* | 10 | Neo | FLMNH | 20204 |
| *Podocarpus lambertii* | 10 | Neo | STRI-CTPA | 898 |
| *Podocarpus lawrencei* | 10 | Aust | STRI - Graham | 19839 |
| *Podocarpus lucienii* | 10 | Aust | STRI - Graham | 22391 |
| *Podocarpus macrophyllus* | 10 | Palearc | STRI - Graham | 63458 |
| *Podocarpus matudae* | 11 | Neo | STRI - Graham | 10584 |
| *Podocarpus milanjianus* | 10 | Afr | Utrecht | NA |
| *Podocarpus neriifolius* | 10 | Aust/Indomalay | STRI - Graham | 19844 |
| *Podocarpus nivalis* | 10 | Aust | STRI - Graham | 12961 |
| *Podocarpus novae-caledoniae* | 11 | Aust | STRI - Graham | 22392 |
| *Podocarpus nubigenus* | 11 | Neo | STRI - CTPA | 1392 |
| *Podocarpus oleifolius* | 11 | Neo | STRI - CTPA | 1393 |
| *Podocarpus parlatorei* | 9 | Neo | STRI - Graham | 8345; 8349 |
| *Podocarpus pilgeri* | 10 | Indomalay | STRI - Graham | 19845 |
| *Podocarpus polystachyus* | 5 | Aust/Indomalay | Utrecht | 19849 |
| *Podocarpus salignus* | 10 | Neo | STRI - Graham | 8346 |
| *Podocarpus sellowii* | 10 | Neo | STRI - CTPA | 1395 |
| *Podocarpus spinulosus* | 12 | Aust | STRI - Graham | 19841 |
| *Podocarpus sprucei* | 12 | Neo | FLMNH | 20207 |
| *Podocarpus steyermarkii* | 10 | Neo | STRI-CTPA | 901 |
| *Podocarpus sylvestris* | 10 | Aust | STRI - Graham | 19833 |
| *Podocarpus tepuiensis* | 10 | Neo | STRI-CTPA | 902 |
| *Podocarpus totara* | 10 | Aust | STRI - Graham | 19843 |
| *Podocarpus urbanii* | 10 | Neo | STRI - Graham | 427685 |

STRI = Smithsonian Tropical Research Institute; CPTA = Center for Tropical Paleoecology and Archaeology; FLMNH = Florida Museum of Natural History

Table S2. *Podocarpidites* fossil specimen slide and sample details. Ages were calculated following the maximum likelihood-based biostratigraphic method described in Punyasena et al. (2012). The specimens include pollen types previously described in Jaramillo et al. (2014), Martínez *et al.*, (2020), and Carvalho et al. (2021). Images from Punyasena et al. (2025).

| Specimen ID | Specimen | Country | Latitude | Longitude | Age range (Ma) |
|---|---|---|---|---|---|
| 1 | Panama (3.6 Ma) | Panama | 9.087969 | -81.536114 | 3.55 |
| 2 | Panama (4.3 Ma) | Panama | 9.17641667 | -82.058 | 4.25 |
| 3 | Brazil 1 (5.3 Ma) | Brazil | -5.3269398 | -71.033775 | 5.3-2.6 |
| 4 | Brazil 2 (5.3 Ma) | Brazil | -5.3269398 | -71.033775 | 5.3-2.6 |
| 5 | Brazil 3 (5.3 Ma) | Brazil | -5.3269398 | -71.033775 | 5.3-2.6 |
| 6 | Brazil 4 (5.3 Ma) | Brazil | -5.3269398 | -71.033775 | 5.3-2.6 |
| 7 | Brazil 5 (5.3 Ma) | Brazil | -5.3269398 | -71.033775 | 5.3-2.6 |
| 8 | Brazil 6 (5.3 Ma) | Brazil | -5.3269398 | -71.033775 | 5.3-2.6 |
| 9 | Brazil 7 (5.3 Ma) | Brazil | -5.3269398 | -71.033775 | 5.3-2.6 |
| 10 | Venezuela 1 (5.5 Ma) | Venezuela | 11.4233476 | -69.642345 | 5.4903 |
| 11 | Venezuela 2 (5.5 Ma) | Venezuela | 11.4233476 | -69.642345 | 5.4903 |
| 12 | Panama (10.2 Ma) | Panama | 8.13188889 | -77.658 | 10.15 |
| 13 | Venezuela (10.6 Ma) | Venezuela | 11.1537085 | -70.270515 | 10.606 |
| 14 | Venezuela (11.8 Ma) | Venezuela | 11.1537085 | -70.270515 | 11.8223 |
| 15 | Peru 1 (12 Ma) | Peru | -14.6663 | -71.2832 | 12-10 |
| 16 | Peru 2 (12 Ma) | Peru | -14.7276 | -71.2674 | 12-10 |
| 17 | Peru 3 (12 Ma) | Peru | -14.6624 | -71.2758 | 12-10 |
| 18 | Peru 4 (12 Ma) | Peru | -14.6624 | -71.2758 | 12-10 |
| 19 | Brazil (13.5 Ma) | Brazil | -4.0374589 | -69.472285 | 13.51-13.25 |
| 20 | Venezuela (13.9 Ma) | Venezuela | 11.1831 | -70.05 | 13.928 |
| 21 | Brazil (14.7 Ma) | Brazil | -3.5238065 | -68.850568 | 14.64-14.02 |
| 22 | Brazil (16 Ma) | Brazil | -4.0374589 | -69.472285 | 16.03-16 |
| 23 | Brazil 1 (16.1 Ma) | Brazil | -4.8833333 | -70.15 | 16.09-16.04 |
| 24 | Brazil 2 (16.1 Ma) | Brazil | -4.0374589 | -69.472285 | 16.09-16.07 |
| 25 | Brazil 1 (16.3 Ma) | Brazil | -4.8833333 | -70.15 | 16.27-16.13 |
| 26 | Brazil 2 (16.3 Ma) | Brazil | -4.0374589 | -69.472285 | 16.27-16.21 |
| 27 | Brazil 3 (16.3 Ma) | Brazil | -5.2690884 | -71.551079 | 16.27-16.21 |
| 28 | Brazil (16.5 Ma) | Brazil | -5.2690884 | -71.551079 | 16.48-16.41 |
| 29 | Brazil 1 (23.3 Ma) | Brazil | -3.4312855 | -67.541133 | 23.3-5.3 |
| 30 | Brazil 2 (23.3 Ma) | Brazil | -3.4312855 | -67.541133 | 23.3-5.3 |
| 31 | Colombia (67.2 Ma) | Colombia | 6.55 | -73.7 | 67.2-67-1 |
